# Characterizing the interplay of rubisco and nitrogenase enzymes in anaerobic-photoheterotrophically grown *Rhodopseudomonas palustris* CGA009 through a genome-scale metabolic and expression model

**DOI:** 10.1101/2022.03.03.482919

**Authors:** Niaz Bahar Chowdhury, Adil Alsiyabi, Rajib Saha

## Abstract

*Rhodopseudomonas palustris* CGA009 (*R. palustris*) is a gram negative purple non-sulfur bacteria that grows phototrophically or chemotrophically by fixing or catabolizing a wide array of substrates including lignin breakdown products (e.g., *p*-coumarate) for its carbon and nitrogen requirements. It can grow aerobically or anaerobically and can use light, inorganic, and organic compounds for energy production. Due to its ability to convert different carbon sources into useful products in anaerobic mode, this study, for the first time, reconstructed a metabolic and expression (ME-) model of *R. palustris* to investigate its anaerobic-photoheterotrophic growth. Unlike metabolic (M-) models, ME-models include transcription and translation reactions along with macromolecules synthesis and couple these reactions with growth rate. This unique feature of the ME-model led to nonlinear growth curve predictions which matched closely with experimental growth rate data. At the theoretical maximum growth rate, the ME-model suggested a diminishing rate of carbon fixation and predicted malate dehydrogenase and glycerol-3 phosphate dehydrogenase as alternate electron sinks. Moreover, the ME-model also identified ferredoxin as a key regulator in distributing electrons between major redox balancing pathways. Since ME-models include turnover rate for each metabolic reaction, it was used to successfully capture experimentally observed temperature regulation of different nitrogenases. Overall, these unique features of the ME-model demonstrated the influence of nitrogenases and rubiscos on *R. palustris* growth and predicted a key regulator in distributing electrons between major redox balancing pathways, thus establishing a platform for *in silico* investigation of *R. palustris* metabolism from a multi-omics perspective.

**IMPORTANCE:** In this work, we reconstructed the first ME-model for a purple non-sulfur bacterium (PNSB). Using the ME-model, different aspects of *R. palustris* metabolism were examined. First, the ME-model was used to analyze how reducing power entering the *R. palustris* cell through organic carbon sources gets partitioned into biomass, carbon dioxide fixation, and nitrogen fixation. Furthermore, the ME-model predicted electron flux through ferredoxin as a major bottleneck in distributing electrons to nitrogenase enzymes. Next, the ME-model characterized different nitrogenase enzymes and successfully recapitulated experimentally observed temperature regulations of those enzymes. Identifying the bottleneck responsible for transferring electron to nitrogenase enzymes and recapitulating the temperature regulation of different nitrogenase enzymes can have profound implications in metabolic engineering, such as hydrogen production from *R. palustris*. Another interesting application of this ME-model can be to take advantage of its redox balancing strategy to gain understanding on regulatory mechanism of biodegradable plastic production precursors, such as polyhydroxybutyrate (PHB).

## INTRODUCTION

*R. palustris* is an alphaproteobacterium which can grow in diverse metabolic modes such as phototrophic or chemotrophic growth. Besides, it can grow under aerobic or anaerobic conditions by using light and organic (e.g., lignin breakdown products) or inorganic compounds as a source of ATP generation (1,2). Using these metabolic versatilities, *R. palustris* has emerged as a potential biotechnological platform for bioremediation (3–5), bioplastics production (6,7), bioelectricity generation (8,9), wastewater treatment (10–12), and hydrogen production (13–17). Furthermore, *R. palustris* is the only known bacteria to encode all three known nitrogenase enzymes (2) besides *Azotobacter vinelandii* (*A. vinelandii*) (18). *R. palustris* also encodes both form I and form II of rubisco. These unique features make *R. palustris* an ideal microorganism to be considered as a biotechnological chassis for further metabolic engineering (7). Because of these unique features, *R. palustris* has a highly connected metabolic network which requires a systems-level investigation for better understanding.

One widely accepted systems level investigation tool is the stoichiometric constrain-based M-model (19). Initial efforts of reconstructing M-models of purple non-sulfur bacteria (PNSB) were limited to the specific metabolic pathways of interest, such as central carbon metabolism (20), and electron transport chain (21). However, those pathway specific M-models did not have wider resolution to capture overall metabolic landscape of PNSBs. To overcome that, comprehensive M-models were reconstructed for PNSB strains including *Rhodobacter sphaeroides* (22) and *R. palustris* (23). Recently we further refined the *R. palustris* M-model by integrating the annotated metabolic pathways for lignin monomer degradation and validated it by using the experimental data on gene essentiality and metabolic flux analysis for growth under different carbon sources (24). Although, these M-models were useful to study different metabolic features of PNSB, the inherent lacking of quantitative characterization of macromolecular machinery synthesis (MMS) could be problematic and may lead to incorrect predictions of biological scenarios, such as inaccurate reaction flux and multiple equivalent cellular phenotypic states (25,26). These inaccuracies can lead to an erroneous understanding of overall metabolic and regulatory features of an organism and can negatively impact the design-build-test-learn cycle for metabolic engineering application.

One of the ways to overcome this is the metabolic-expression (ME) modeling approach. ME-model is a resource allocation based model that includes not only the stoichiometric metabolic reactions, but also quantitative MMS information (27). As input, ME-models require the conditions of a steady-state environment and can then output predictions for maximum growth rate, substrate uptake, byproduct secretion, metabolic fluxes, gene expression levels, and protein expression (27). ME-model utilizes a growth optimization function along with coupling constraints that tie flux to transcriptional and translational reactions in the model. These constraints are functions of the growth rate. By including these constraints, ME-models set limitations on fluxes based on transcription as well as translation reactions. Thus far, ME-models were developed only for a few organisms. These models were used to accurately predict cellular composition and gene expression of *Thermotoga maritima* (*T. maritima*) (28), fermentation profile of *Clostridium ljungdahlii* (*C. ljungdahlii*) (29), overflow metabolism of *Saccharomyces cerevisiae* (*S. cerevisiae*) (30), and multi-scale phenotype, enzyme abundance, and acid stress of *Escherichia coli* (*E. coli*) (31–33). An ME-model for *R. palustris* can also be very useful in answering fundamental biological questions, such as growth profiling, isozyme expression prediction, regulation on electron distribution between competing metabolic modules, and temperature regulation of different enzymes.

In this work, the first ever ME-model was reconstructed for *R. palustris*. The ME-model was able to satisfactorily recapitulate the experimental transcriptomics and proteomics observation from literature (34). Then acetate, succinate, butyrate, and *p*-coumarate were used as carbon sources to characterize the growth profile of *R. palustris* which closely matched with experimental growth rate data. In addition, it predicted a diminishing rate of carbon fixation at the theoretical maximum growth rate and consequently predicted malate dehydrogenase and glycerol-3 phosphate dehydrogenase as alternate electron sinks. Furthermore, the ME-model identified ferredoxin as a key regulator in distributing electrons between major redox balancing pathways, such as carbon and nitrogen fixation. Later, the modeling framework was able to capture experimentally observed temperature regulation of different nitrogenase enzymes, by varying turnover rate of nitrogen fixation reactions. Overall, this modeling approach demonstrated a bottom-up systems-biology approach that can be used to predict and analyze cellular physiology of *R. palustris*, thereby providing an opportunity to generate experimentally testable hypotheses.

## RESULTS AND DISCUSSIONS

### Metabolic and expression model development

To reconstruct the ME-model, our previously reconstructed M-model of *R. palustris, iRpa940* (24), was used as a template for the metabolic transformations. To reconstruct the ME-model, gene-protein-reaction (GPR) relationships for all the reactions, specially nitrogen fixation (catalyzed by Mo-, V-, and Fe-Nase) and carbon fixation (catalyzed by rubisco form I and form II) reactions, were manually curated from the complete genome sequence of *R. palustris* (2). Transcription and translation reactions were added for reactions for which GPR relationships are available. Reactions for which GPR associations are not available, it was assumed that an average bacterial enzyme with 31.09 *kDa* molecular weight (35) catalyzed each individual reaction. Overall, the ME-model contains 1398 reactions, 1483 metabolites, and 751 genes. FIG 1 demonstrates the workflow of the ME-model reconstruction.

**FIG 1.**
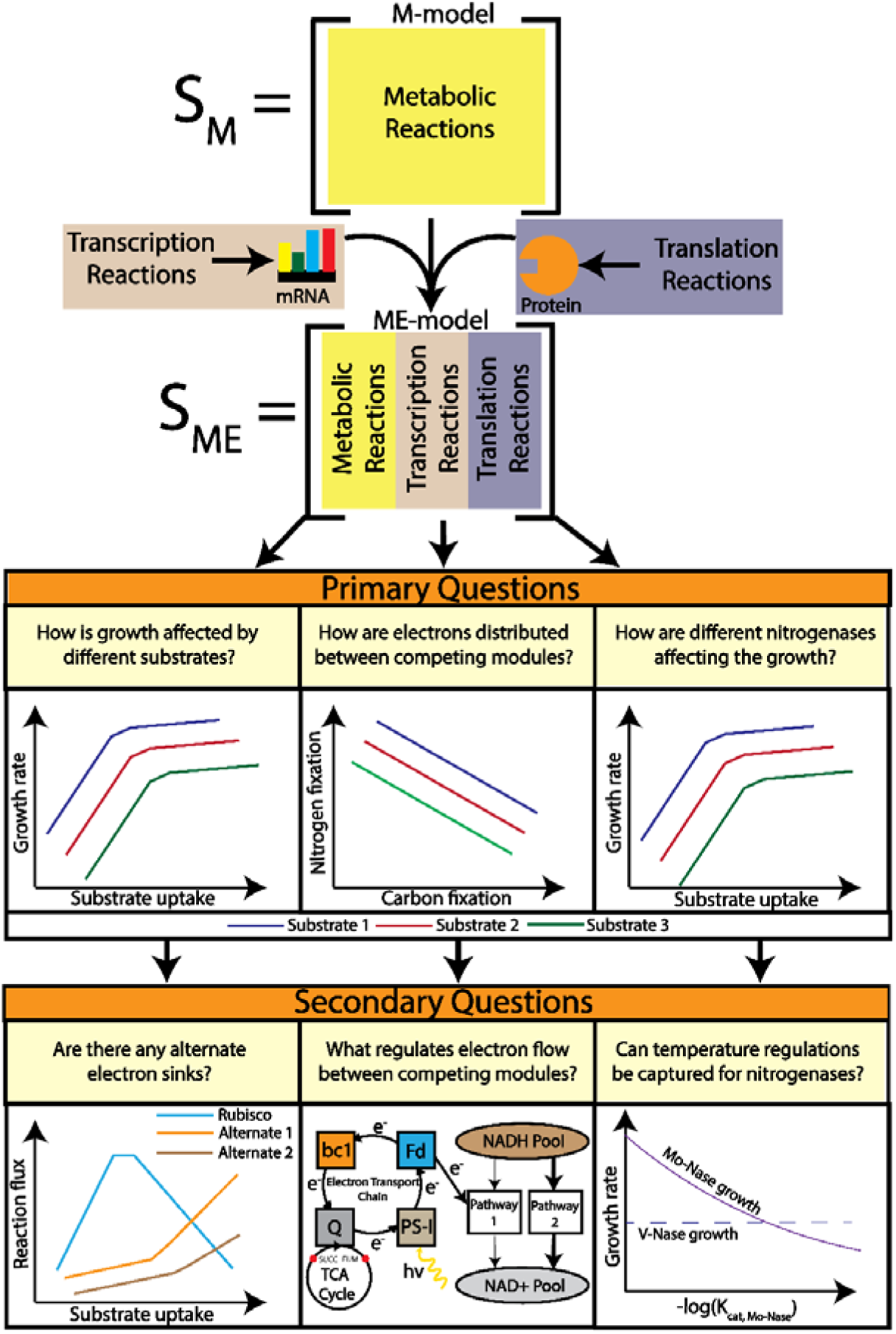
Workflow followed to reconstruct the ME-model from a previously published M-model of *R. palustris*. Transcription and translation reactions were added on top of the metabolic reactions to come up with ME-modeling framework. The ME-modeling framework was used to characterize growth rate profiling, competing metabolic modules, and nitrogenase enzyme activity. From these characterizations, inferences regarding alternate redox balancing, ferredoxin regulation, and temperature regulation of nitrogenase enzymes were gathered.

In *R. palustris*, form I rubisco (L_8_S_8_) is comprised of eight large subunits (L_8_) and eight small subunits (S_8_) (36) and encoded by two genes, *rpa1559* and *rpa1560* (2). On the other hand, form II rubisco (L_2_) is comprised of two large subunits both encoded by *rpa4641* (2). Between the two forms of rubisco, form I has a higher molecular weight compared to form II (37,38) and therefore requires more carbon investment to synthesize. As rubisco is one of the most abundant enzymes in nature (39), the kinetics of this enzyme have been determined for multiple organisms (36,40). For different rubisco enzymes, it was shown that although form I has higher molecular weight and more carbon investment cost, form II has higher catalytic turnover rate (*k*_*cat*_) per active site compared to form I (36). Evolutionary selection has played a major role in this counterintuitive observation (41,42). Early in the earth’s history, the concentration of carbon dioxide was higher in the atmosphere and as a result form II rubisco evolved with a lower selectivity and higher *k*_*cat*_ for carbon dioxide (36). With increasing amounts of oxygen in earth’s atmosphere, form I evolved with a much higher selectivity for carbon dioxide but with a lower *k*_*cat*_ (36). Since *k*_*cat*_ values for *R. palustris* are not available, to account for these evolutionary selections, the *k*_*cat*_ values were set to 3.7 *s*^-1^*active site*^−1^ (form I) and 6.6 *s*^−1^*active site*^−1^ (form II) based on the measurements from other phylogenetically close (43) PNSB strains (*Rhodobacter capsulatus* (40) and *Rhodospirillum rubrum* (36), respectively).

For the three nitrogenase isozymes, each enzyme is encoded by a series of genes (2) (Mo-Nase by *rpa4602* - *rpa4633*, V-Nase by *rpa1370* - *rpa1380*, and Fe-Nase by *rpa1435* - *rpa1439*). Unlike rubisco, *k*_*cat*_ values of different nitrogenase are not available for *R. palustris* or any other PNSBs. Therefore, the calculated surface accessible surface area (SASA) of each nitrogenase enzyme was used to normalize the mean *k*_*cat*_ value from *E. coli*, as discussed in literature (31) (see materials and methods section). These normalized *k*_*cat*_ values were used to define three independent nitrogen fixation reactions.

Both of the above mentioned enzymes, nitrogenase and rubisco, play a pertinent role in maintaining the cellular redox balance during the photoheterotrophic growth of *R. palustris* by regenerating oxidized cofactors (44). When the ME-model was used to simulate the photoheterotrophic growth of *R. palustris*, among three different nitrogenase enzymes, it predicted the expression of Mo-Nase only, which is consistent with literature (45,46). For the same photoheterotrophic growth conditions, between two different forms of rubisco enzymes, the model predicted only the expression of form II rubisco. Although expression of only rubisco form II was expected based on its lower carbon cost and higher efficiency, literature evidence suggested a co-expression of both forms of rubisco during the photoheterotrophic growth of *R. palustris* (47). The same work suggested that rubisco form I is responsible for providing cellular carbon and dominates under carbon dioxide limiting conditions, whereas rubisco form II balances the intracellular redox potential under carbon and electron abundant conditions (47). In addition, it was also found that expression of the *cbb* operons (responsible for coding both forms of rubisco) during phototrophic growth is highly dependent on the cellular carbon dioxide level (47). To incorporate these findings, a constraint was added to the ME-model to co-express both forms of rubisco based on the total carbon dioxide produced by *R. palustris* during photoheterotrophic growth (see materials and methods section).

### Model Validation using experimental transcriptomics and proteomics data

To validate the prediction accuracy of the model, experimental transcriptomics and proteomics data were used to qualitatively verify whether the model can predict the direction of these experimental fold changes in different conditions. A previous study, which characterized the anaerobic growth of *R. palustris* by comparing the transcriptomic and proteomic profiles of cultures grown in the presence of *p*-coumarate and succinate as sole carbon source, was used for the validation study (34). The study tested fold change of 4810 genes for *p*-coumarate catabolism considering succinate catabolism as the baseline condition. The transcriptomic analysis resulted in 369 differentially expressed genes, among which 61 were metabolic genes. Similarly, proteomic analysis resulted in 341 differentially expressed proteins, among which 67 can act as enzymes. In both transcriptomics and proteomics data sets, non-metabolic genes/proteins have functions such as signaling, chromosomal replication, and circadian rhythm. (see supplemental material Table S1 for more details).

To generate both gene and protein expression information for the same two conditions of the above-mentioned study (34), the ME-model was simulated for two points where total rubisco flux was maximal for the *p*-coumarate and succinate uptake, respectively. It was previously reported (44) that carbon fixation is required to maintain redox balance in *R. palustris*. Therefore, higher growth rate is associated with higher reduced cofactor production, leading to higher rates of carbon fixation. As a result, the decreasing carbon fixation flux with increasing growth (FIG 2) is a theoretical feature predicted by the ME-model. All the experimentally observed and differentially expressed genes and proteins are available in the model. However, for reactions catalyzed by multiple isozymes, the ME-model only predicted the most efficient isozyme based on the *k*_*cat*_ and molecular weight. As a result, out of these 61 metabolic genes and 67 metabolic enzymes, 23 genes and 34 enzymes were expressed in the model.

**FIG 2.**
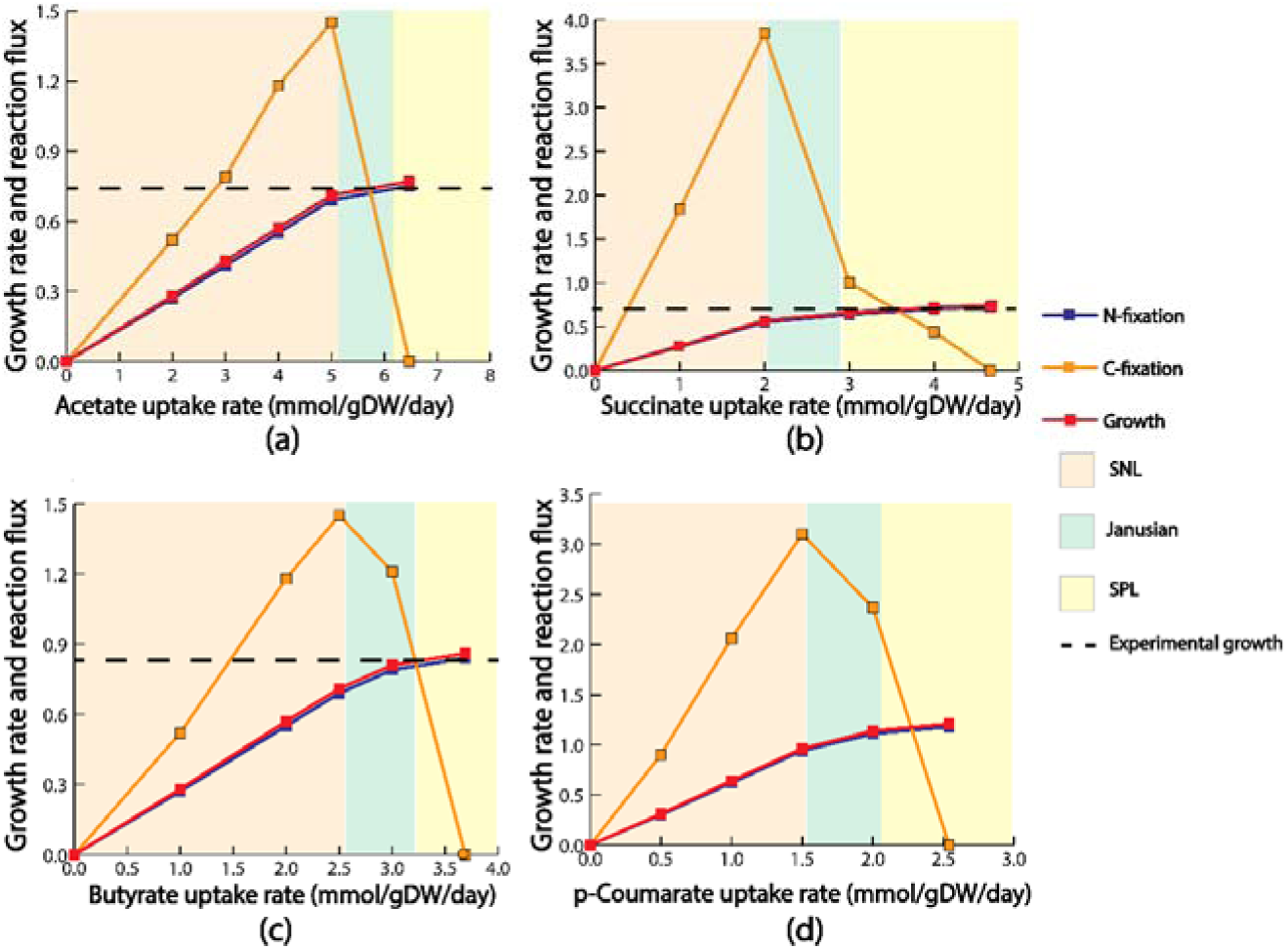
Strictly Nutrient-Limited (SNL), Janusian, and Strictly Proteome-Limited (SPL) regions for (a) acetate (b) succinate (c) butyrate and (d) *p*-coumarate. The growth rate with respect to different substrate uptakes follows a non-linear pattern. Flux through nitrogen fixation reaction also follows the similar pattern to growth rate. Carbon fixation reached a peak in the Janusian region and then diminished in the theoretical maximal growth.

As part of the transcriptomics data validation, out of 23 genes, the ME-model was able to predict correct gene expression fold change for 21 genes. The model could not predict the downward fold change of 3-oxoacyl-acyl carrier protein reductase (*rpa3304*) and the 50S ribosomal protein (*rpa0918*). *rpa3304* is one of the genes to convert malonyl-CoA to biotin (48). Biotin is a part of *R. palustris* cell membrane and from FIG 2 it can be seen that *p*-coumarate supports more growth than succinate. Thus, the ME-model predicted an upward fold change of *rpa3304* for *p*-coumarate catabolism compared to succinate catabolism. Composition of biotin in cell membrane may be different in different conditions. However, in the ME-model, only protein and nucleotide compositions change with different conditions, while those of cell wall components remain constant (49). This may have caused the mismatch. For incorrect fold change prediction of 50S ribosomal protein, missing reactions, the lack of regulatory mechanisms, and inaccurate *k*_*cat*_ data may have played a role.

For proteomics data validation, out of 34 enzymes, the ME-model was able to correctly predict the fold change for 21 enzymes. The ME-model could not correctly predict the downward fold change of 13 different enzymes (see supplemental material Table S1 for more details). These enzymes are mainly associated with purine and pyrimidine metabolism, fatty acid metabolism, and lipopolysaccharide metabolism. These pathways are closely associated with the *R. palustris* biomass growth. As *p*-coumarate supports more growth than succinate, the ME-model allocated more proteins for these pathways to sustain the biomass growth. There may be unannotated alternate metabolic pathways with less enzyme investment for producing purine, pyrimidine, fatty acid, and lipopolysaccharide when *p*-coumarate is utilized as the carbon source, thus causing these discrepancies. As ME-model maximizes the biomass growth rate, such incorrect prediction can be considered as an inherent weakness of the ME-model.

Overall, despite these incorrect fold change predictions, the ME-model was able to satisfactorily recapitulate the aggregate experimental transcriptomics and proteomics observations with 91% and 62% accuracy, respectively (see materials and methods section for accuracy calculation). The details of experimental and model predictions can be found in the supplemental material Table S1.

### Growth rate vs. substrate uptake and alternate redox balancing strategies

Upon the validation with available gene expression and protein abundance data, the model was used to examine how growth, carbon fixation, and nitrogen fixation rates varied with different substrate uptake rate. The goal of this analysis was to investigate how reducing power entering the cell through organic carbon sources gets partitioned into biomass, carbon dioxide fixation, and nitrogen fixation. To perform the analysis, acetate, succinate, butyrate, and *p*-coumarate were used as substrates. Previous studies have shown that photoheterotrophic growth of *R. palustris* on acetate, succinate, and butyrate is associated with increasing cellular redox stress based on the oxidation state of different substrates (50). Hence, these substrates were chosen as they cover a wide range of oxidation states. Here succinate (+0.5) and acetate (0) have higher oxidation states compared to *R. palustris’* biomass (−0.13) (45), whereas butyrate (−1) and *p*-coumarate (−0.22) have lower oxidation states (45).

In the ME-model, growth rate is a nonlinear function of substrate uptake rate and eventually reaches a theoretical maximum growth rate (FIG 2). This behavior is consistent with known microbial empirical growth models such as Monod growth kinetics (51) and microbial slow growth kinetics (52). Previous work has suggested three distinct growth regions as a function of substrate uptake rate; Strictly Nutrient-Limited (SNL), Janusian, and Strictly Proteome-Limited (SPL) (31). Growth in the SNL region depends heavily on nutrient uptake and adding more nutrient results in more growth. In this region, the relationship between growth rate and substrate uptake is similar to the prediction made from M-models. Contrary to the SNL region, growth in the SPL region (also known as nutrient excess condition) is limited by physiological constraint of protein production and catalysis. Janusian growth is the region where a transition from SNL to SPL takes place. A recent experimental study (45) had characterized the growth of wild-type (WT) *R. palustris* for acetate, succinate, and butyrate, respectively, under nitrogen-fixing conditions. Table 2 compares between experimentally observed growth rates and those predicted by the model. The growth rate and order predicted by the ME-model for succinate, acetate, and butyrate closely followed the experimental growth rate and order. Compared to other substrates, the ME-model predicted a significantly higher growth rate on *p*-coumarate. One of our previous works (7), which experimentally examined different strategies for PHB production under non-nitrogen fixing condition, also showed a significantly higher growth on p-coumarate comparing to butyrate and acetate. It was previously reported (7) that, *p*-coumarate consumption lead to more ATP production compared to acetate, succinate, and butyrate and thus was able to support more growth.

**Table 2:** Normalized growth rate for different substrate uptakes.

| Substrate | Experimentally<br>observed growth rate<br>$day^{-1}$ | Growth rate from the<br>ME-model ($day^{-1}$) | Substrate Uptake for<br>experimental growth<br>rate from model<br>($mmol. gDW^{-1}. day^{-1}$) |
| --- | --- | --- | --- |
| Succinate | 0.70 | 0.74 | 4.66 |
| Acetate | 0.74 | 0.77 | 6.47 |
| Butyrate | 0.82 | 0.86 | 3.69 |
| *p*-Coumarate | - | 1.21 | 2.54 |

Theoretical growth rates predicted by the ME-model were slightly higher compared to the experimental growth rates for all tested substrates (6% for succinate, 5% for butyrate, and 4% for acetate). It was expected as the cell has many more layers of physiological regulations, such as signaling pathways, allosteric regulation, and polymorphism, which were not captured in the ME-modeling framework. Overall, growth rate comparison between the ME-model prediction and experimental study reveals that, like *E. coli* (31), optimum resource allocation dictates metabolic activities for *R. palustris*. Supplemental material Table S2 records all the theoretical maximum growth rates for different amount of substrate uptakes.

After characterizing the growth rate with different substrate uptakes, the ME-model was used to characterize nitrogen and carbon fixation rates as a function substrate uptake. For nitrogen fixation, the reaction’s activity followed a similar trajectory as growth vs. substrate uptake (FIG 2). Different studies have shown that during WT photoheterotrophic growth, among three different nitrogenase (Mo-, V-, and Fe-Nase) isozymes encoded in *R. palustris’* genome, Mo-Nase is exclusively expressed (45,46). *A. vinelandii* which has three different nitrogenases also exclusively express the Mo-Nase in the WT (53). The ME-model predicted exclusive expression of Mo-Nase during growth on all four carbon sources. Expression of nitrogenase may be dictated by its ATP requirements, as Mo-Nase requires the least amount of ATP among three nitrogenases. In addition, the temperature of the assay plays a role in the expression of different nitrogenases as discussed later.

Next, carbon fixation was also characterized with respect to substrate uptake. Unlike nitrogenase, which closely followed the trajectory of the growth rate, carbon fixation reached a peak flux at the start of the Janusian region. In the SPL region, when growth is proteome limited, *R. palustris* optimized protein production to sustain the growing biomass demand. As the cell approaches the theoretical maximal growth, more ribose-5 phosphate is needed to sustain the increasing demand of nucleotides and lipopolysaccharides. To meet that demand at the theoretical maximum growth, the ME-model predicted that *R. palustris* decreases the expression of phosphoribulokinase (*rpa4645*) and redirects flux towards ribose-5 phosphate production (FIG 3).

**FIG 3.**
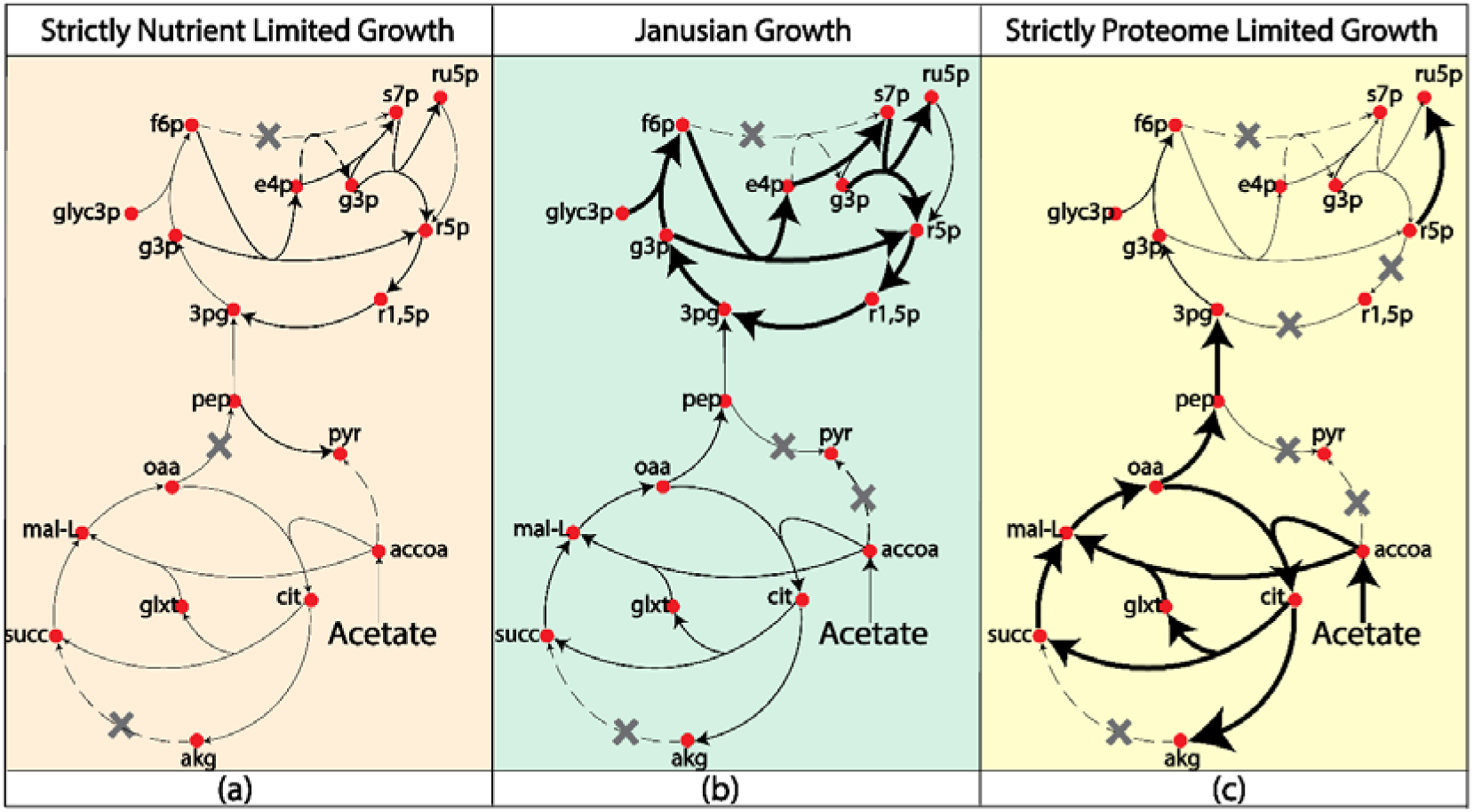
Metabolic activities in the (a) strictly nutrient limited growth (SNL), (b) Janusian growth, and (c) strictly proteome limited growth (SPL). In the theoretical maximum growth, at SPL region, flux through carbon fixation diminished and reaction flux from ribulose-5 phosphate to ribose-5 phosphate significantly increased. The increased biomass growth demand can be met by the precursors from the TCA cycle, which showed significant increase in reaction flux comparing to Janusian growth and SNL. Here gray crosses indicate zero rection flux through that reaction.

During photoheterotrophic growth under nitrogen fixing condition, carbon and nitrogen fixation plays a major role in maintaining cellular redox balance. However, in the SPL region, as reaction flux of carbon fixation diminished at the theoretical maximum growth, the ME-model predicted two potential candidates to maintain cellular redox balance: malate dehydrogenase and glycerol-3 phosphate dehydrogenase, in addition to nitrogen fixing reaction. Malate dehydrogenase uses NAD+/NADH as cofactors and is encoded by *rpa0192*. Similarly, glycerol-3 phosphate dehydrogenase uses NAD+/NADH as cofactors and is encoded by *rpa4410*. During the switch from the SNL to the SPL region, at the point where carbon fixation starts to diminish, both malate dehydrogenase and glycerol-3 phosphate dehydrogenase fluxes start to increase (FIG 4). At the theoretical maximum growth, flux through malate dehydrogenase and glycerol-3 phosphate dehydrogenase reached its maximum. Malate dehydrogenase also plays a role in maintaining redox balance in several other gram negative bacteria, such as organisms including *E. coli* (54), and *Corynebacterium glutamicum* (*C. glutamicum*) (55). Glycerol-3 phosphate dehydrogenase is one of the key enzymes in the fatty acid biosynthesis. It was suggested that for photoheterotrophically grown *R. rubrum*, it is possible that other biosynthetic pathways such as fatty acid biosynthesis could offer flexibility contributing to the redox balance (56). In addition, several other organisms such as *S. cerevisiae* (57) and *Kluyveromyces lactis* (*K. lactis*) (58) showed evidence of using glycerol-3 phosphate dehydrogenase to maintain redox balance.

**FIG 4.**
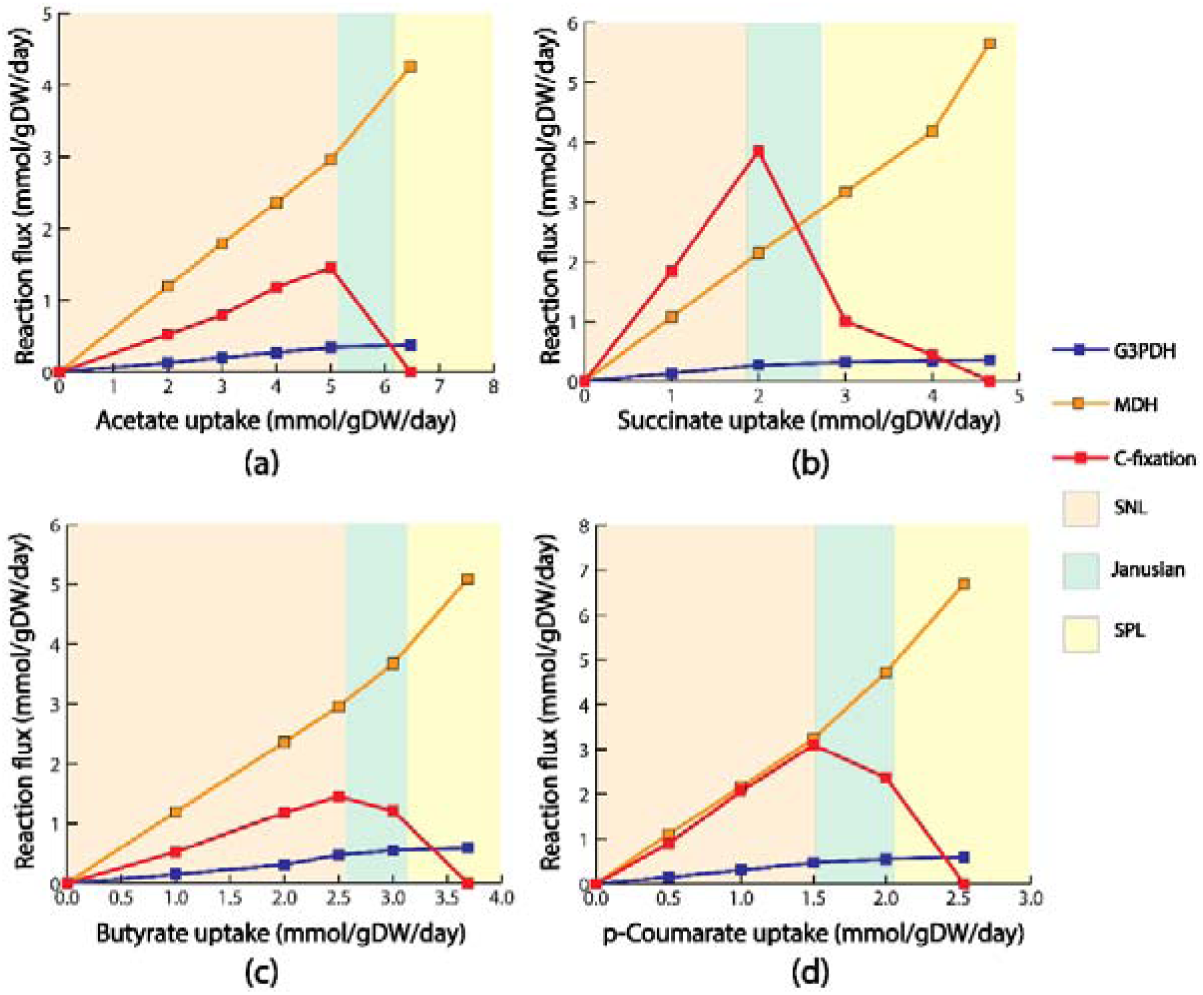
Alternate electron sink for different substrates (a) acetate (b) succinate (c) butyrate and (d). In the Janusian regions, flux through carbon fixation reaction started to diminish. With the diminishing carbon fixation flux, ME-model predicted two alternate electron, malate dehydrogenase and glycerol-3 phosphate dehydrogenase. Reaction flux through these alternate electron sinks reached its peak when flux through carbon fixation completely diminished at the theoretical maximum growth.

### Carbon fixation vs. Nitrogen fixation – competing metabolic modules for redox balance

During photoheterotrophic growth, *R. palustris* performs a cyclic photophosphorylation (2,21) which means that electrons from photosystem I (PSI) get transported through ferredoxin and the *bc*_1_ complex and recycled back to PSI through the oxidation and reduction of quinones (59) (FIG 5). As there are no terminal electron acceptors, this can cause an accumulation of reduced cofactors resulting in impeded growth of the bacterium. To resolve this, *R. palustris* employs various electron acceptors to maintain a cellular redox balance. During photoheterotrophic growth, the redox-balancing mechanism consists primarily of the CBB pathway (44) and nitrogen fixation pathway (60). The nitrogen fixation module becomes active when *R. palustris* is placed in a nitrogen-limiting environment. Experimental studies have suggested a link between carbon and nitrogen fixation that is intimately associated with the control of intracellular redox balance for different PNSBs, such as *R. palustris* (44), *R. capsulatus* (61), *R. sphaeroides* (60,62), and *R. rubrum* (60). However, it is still not properly understood what factors decide the distribution of electrons in these two competing metabolic modules. Here, the ME-model was used to further analyze the metabolic factors deciding the distribution of electron flux between carbon and nitrogen fixation in maintaining cellular redox balance.

**FIG 5.**
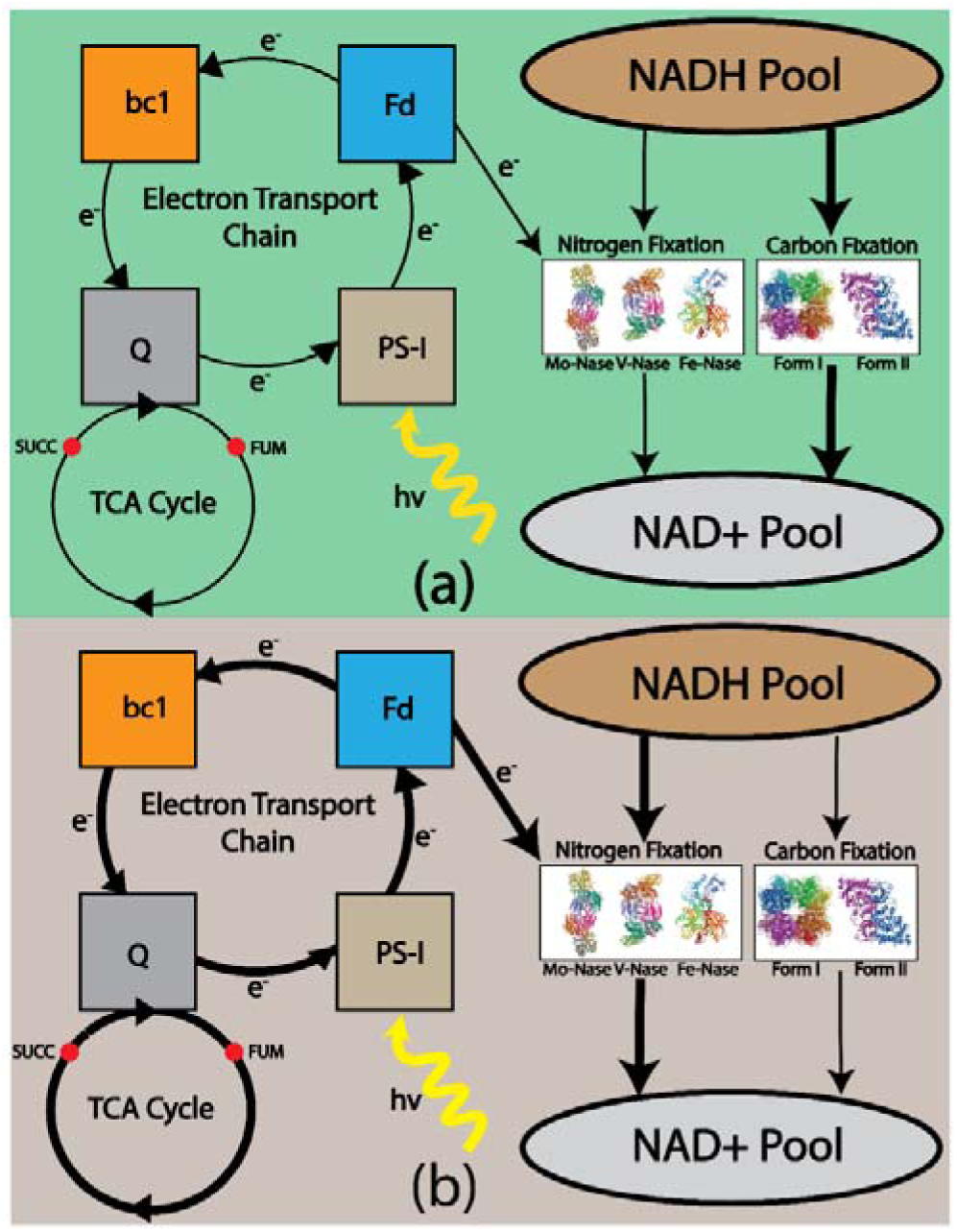
Relation between cyclic photophosphorylation and electron distribution between carbon and nitrogen fixation. (a) Less electron through ferredoxin indicates less flux through nitrogen fixation and more flux through carbon fixation pathway. As a result, NADH will be more oxidized through carbon fixation reaction (b) More electron through ferredoxin indicates more flux through nitrogen fixation and less flux through carbon fixation pathway. As a result NADH will be more oxidized through nitrogen fixation reaction.

To understand the electron distribution, a previous study eliminated rubisco activity in *R. palustris* and found that the rate of nitrogen fixation did not vary significantly (44). As CBB and nitrogen fixation pathways are two major redox balancing mechanisms, when rubisco was eliminated, nitrogen fixation pathway was likely to carry additional flux load to maintain cellular redox balance. As this was not the case in the previous experimental study (44), it was suspected that there exists a metabolic bottleneck preventing additional reaction flux through the nitrogen fixation pathway. FIG 5 shows the cyclic photophosphorylation of *R. palustris*. Here, electrons get transported in a cyclical manner and reduced ferredoxin supplies electrons to nitrogen fixation pathway. Based on the availability of electrons, nitrogen fixation pathway uses reduced cofactors to fix nitrogen. The more electrons supplied by ferredoxin; the more reduced cofactors will be used by nitrogen fixation pathways. Hence, less reduced cofactors will be available for carbon fixation pathway to use. So, electron transport through ferredoxin (ETFD) can be a potential candidate of the previously discussed bottleneck.

In order to explore if ETFD is indeed the hypothesized bottleneck, the biomass growth and substrate uptake rate were kept constant and only flux through carbon fixation reaction was varied for increasing flux of electron transport through the ferredoxin reaction (ETFD). At first, flux through ETFD was fixed to the solution found by the ME-model (indicated by the red line in FIG 6). The flux through the nitrogen fixing reaction remained constant with changing flux through carbon fixation reaction. This finding confirmed the presence of the previously hypothesized bottleneck. Increasing flux through ETFD had varying effects on the rate of nitrogen fixation depending on the utilized carbon substrate. When the reaction flux through ETFD was set to values higher than the ME-model solution (indicated by the yellow and blue lines in FIG 6), a very small change in flux through nitrogenase was noticed for growth on acetate. For the other carbon sources, when the reaction flux through ETFD was set to values higher than the ME-model solution (indicated by the yellow and blue lines in FIG 6), a negative correlation was observed between the carbon and nitrogen fixation reaction flux. When the metabolite pool size (See supplemental material Text S1 for metabolite pool size calculation detail) was calculated for different cofactors, acetate produced less reduced cofactors per unit of substrate uptake compared to other substrates. As, in this case, the fixed nitrogen is the sole source of nitrogen, cell prioritize electron transport to nitrogenase rather than rubisco, whose primary function is to maintain the redox balance in the cell. Thus the relation between carbon and nitrogen fixation is less visible for acetate. However, for succinate, butyrate and *p*-coumarate, more reduced cofactors are produced per unit of substrate uptake. Thus more electrons are available for carbon fixation pathway and the regulation is more visible when ETFD flux is higher for these substrates. These results indicated that reaction flux through ETFD may play a regulatory role in distributing electron flux between carbon and nitrogen fixation.

**FIG 6.**
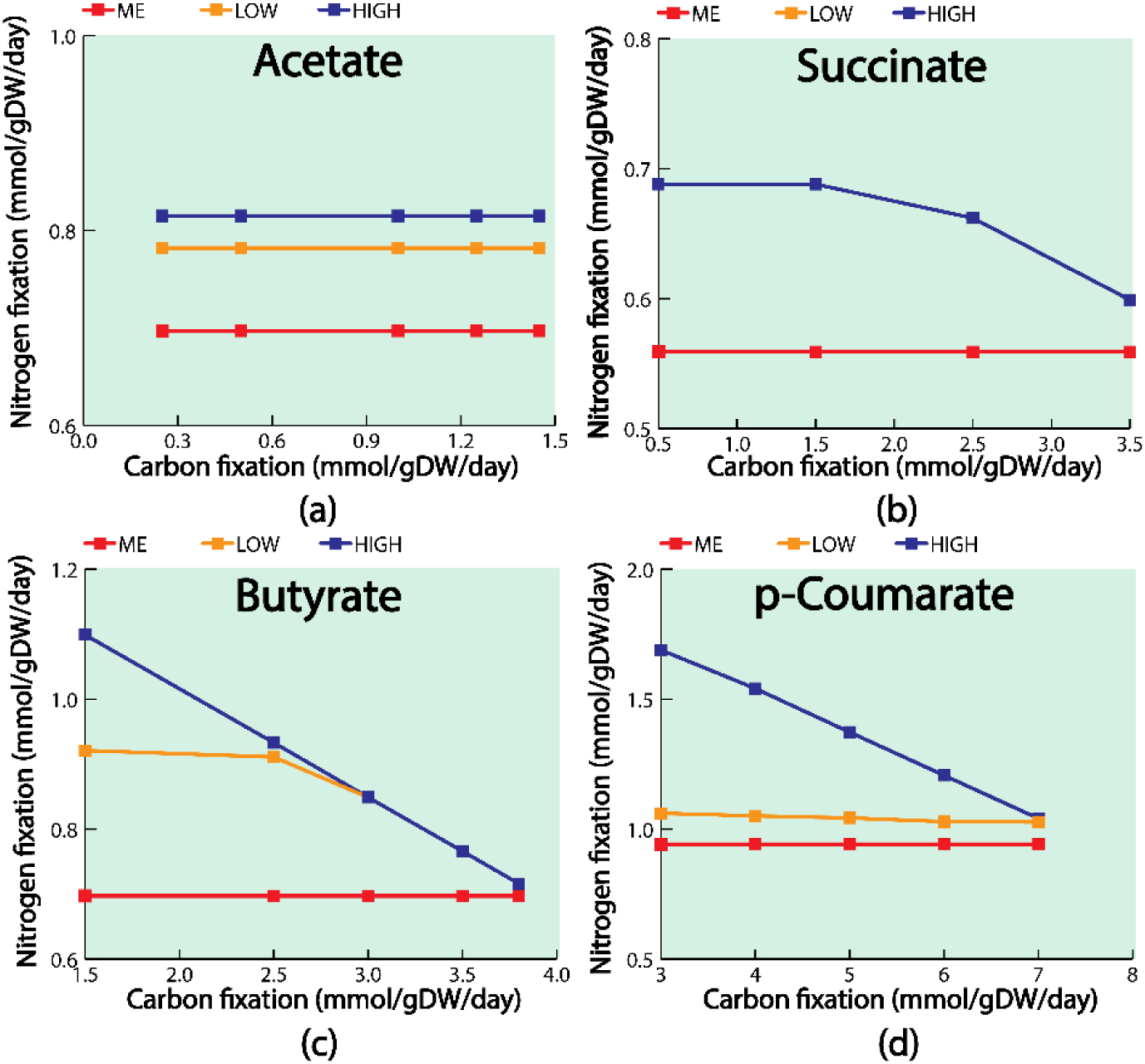
Relation between carbon fixation and nitrogen fixation with different fluxes via electron transport through ferredoxin (ETFD) for different substrates (a) acetate (b) succinate (c) butyrate and (d) *p*-coumarate. Red color lines indicate the relation between carbon fixation and nitrogen fixation when flux through ETFD is set to the solution found from the ME-model. Blue color lines indicate the relation between carbon fixation and nitrogen fixation when ETFD flux is set to a very high value. Yellow color lines indicate the relation between carbon fixation and nitrogen fixation when the ETFD flux values is set between ME and High.

Similar regulation in electron transport between competing metabolic modules, such as respiratory pathways and electron transport, can be observed in model bacteria *E. coli* (63). A highly organized network of overlapping transcriptional regulatory elements regulates flow of electrons by controlling the expression of different genes in *E. coli*, including genes involved in substrates uptake, control of mixed-acid fermentation pathways, and controlling cofactor biosynthesis. Further experimentation is required to establish a similar molecular level mechanism for ETFD regulation of electron distribution in competing pathways of *R. palustris*. The ETFD regulation, hypothesized in this study, can have profound implications in future metabolic engineering efforts of *R. palustris*. Specially, this regulation can be exploited to increase hydrogen production from *R. palustris* to achieve energy sustainability goals.

### Characterization of Mo-, V-, and Fe-Nase nitrogenase enzymes

Since ETFD was postulated to play a regulatory role in distributing electron to the nitrogen fixing pathway, the ME-model was next used to characterize how these electrons were used by different nitrogenase enzymes. First, growth was simulated for the WT *R. palustris* with succinate as the substrate. In this case, only Mo-Nase was expressed and the growth vs. substrate uptake curve (FIG 7 a) followed the pattern identified from the literature (31). Exclusively expressing the Mo-Nase in the WT was also consistent with previous literature findings (45,46). Next, the growth vs. substrate uptake graphs (FIG 7 b, c, and d) were developed for three different mutants of *R. palustris*, each expressing a single nitrogenase isozyme. When the theoretical maximum growths for these mutants were compared with WT, it was found that WT and the Mo-only mutant had the highest growth rate followed by the growth rate of V-only and Fe-only mutants. When compared with the experimental growth rate data from literature (45) for WT, Mo-only, V-only, and Fe-only growths followed a similar pattern as predicted by the ME-model (supplemental material FIG. S1). Theoretically, growth of the WT and -only mutant strains of *R. palustris* can be coupled with the ATP requirement, as Mo-nase requires the least and Fe-nase requires the most amount of ATP for nitrogen fixation.

**FIG 7.**
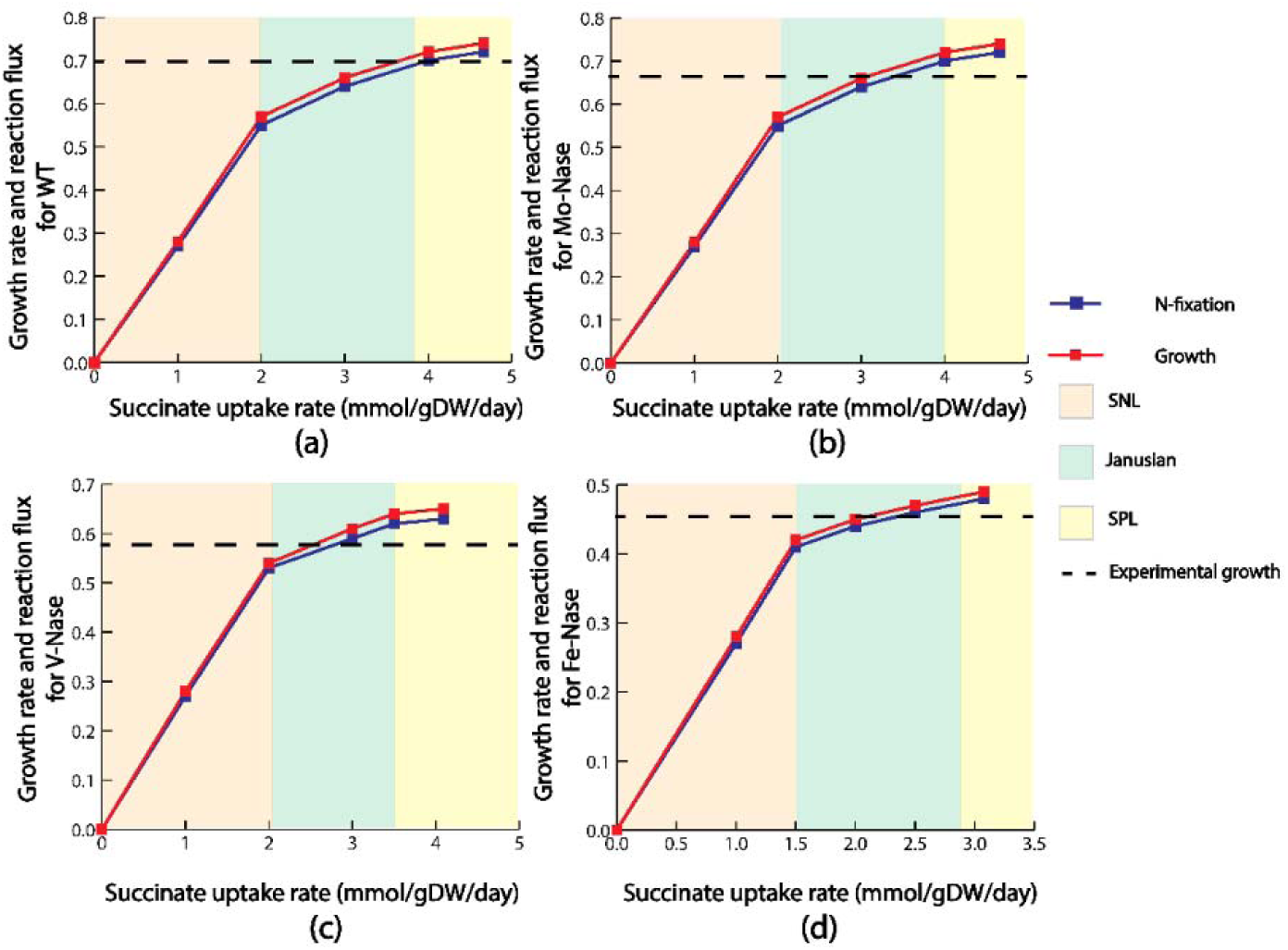
Growth rate and nitrogen fixation rate for (a) WT *R. palustris* (b) Mo-only mutant (c) V-only mutant (d) Fe-only mutant for succinate uptake. For each of the case, growth rate and nitrogen fixation closely follow each other. Dotted line in each of the graph indicates the experimentally observed growth.

Contrary to the pattern observed for succinate uptake, when other carbon sources were used as substrates, V-Nase exhibited higher growth comparing to Mo-Nase, Fe-Nase, and even WT (supplemental material FIG. S2-S4). Previous studies have observed that the Mo-Nase is more sensitive towards decreasing temperature compared to the other isozymes, such as V-Nase (45). FIG 8a qualitatively summarized this idea. Since the experimental values used in that study were generated at 19 °*C*, it is possible that Mo-Nase may have less selectivity towards fixing nitrogen rate than other substrates. The effect of decreasing assay temperature on the activity of nitrogenase is complex. It was reported (64) that for the Mo-Nase of *A. vinelandii*, the rate of nitrogen reduction at 10 °*C* is very low despite continued hydrolysis of ATP. In the case of Mo-Nase of *Klebsiella pneumoniae* (*K. pneumoniae*), decreasing the temperature not only curtails electron flux, but also results in the preferential loss of activity towards nitrogen as a substrate compared with H^+^ or ethyne (C_2_H_2_) (65).

**FIG 8.**
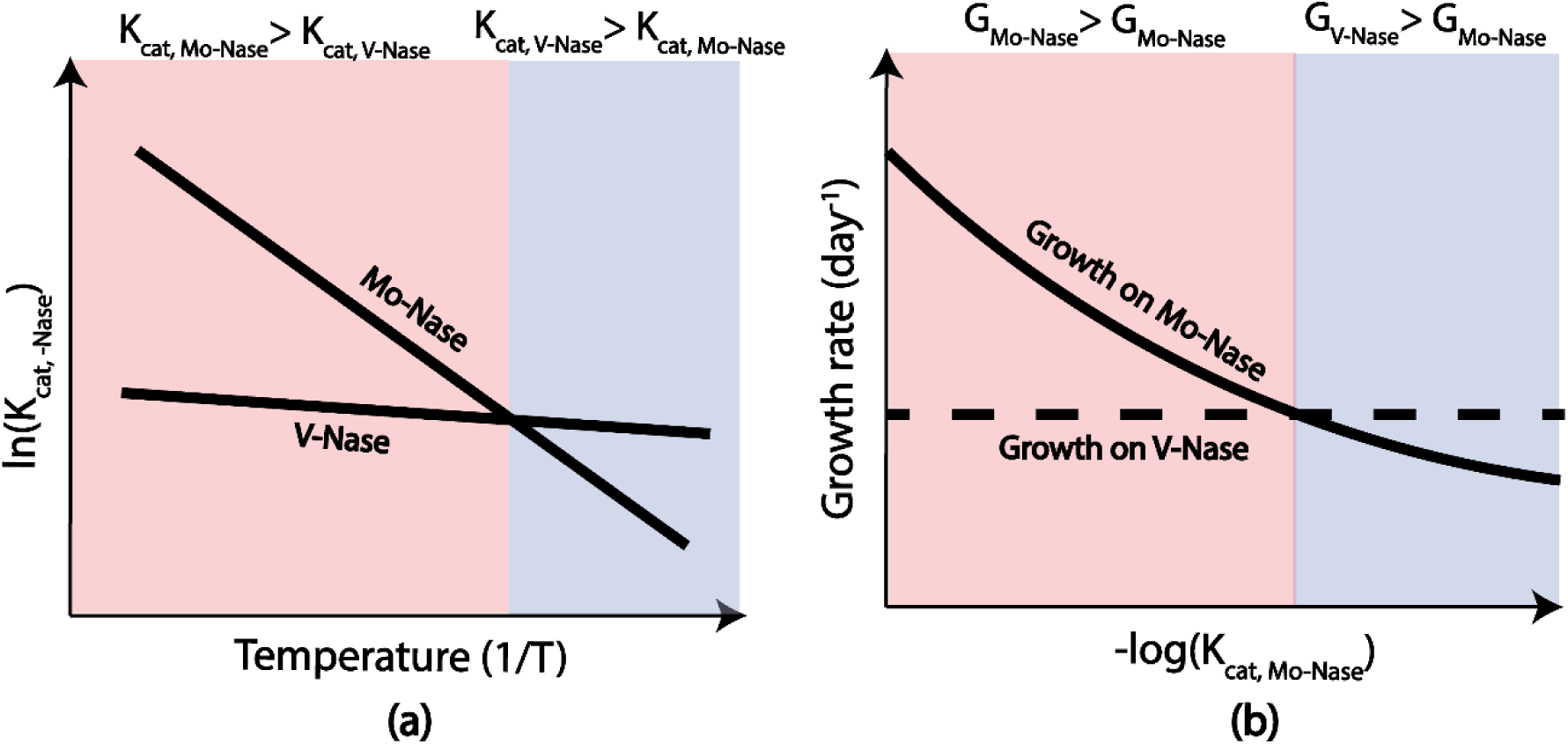
A qualitative representation of temperature regulation of Mo-Nase and V-Nase, and effect of *k*_*cat*_ on the growth of Mo-only and V-only mutant. (a) From literature, it is known that V-Nase has less sensitivity with respect to temperature comparing to the Mo-Nase. The prediction from this ME-model corroborates that finding. (b) As *k*_*cat*_ is a parameter which is a function of temperature, from Arrhenius equation, we know that with reducing temperature, *k*_*cat*_ also reduces. With reducing *k*_*cat*_, at one stage, growth for Mo-only mutant falls below the growth of V-only mutant, capturing the experimentally observed temperature regulation of Mo- and V-Nase.

This modeling framework was further used to investigate the decreased growth rate of Mo-Nase at lower temperature. From Arrhenius equation (66) it is known that turnover rate of an enzyme, *k*_*cat*_, increases exponentially with the increasing temperature. As *k*_*cat*_ is one of the temperature sensitive parameters in this study, *k*_*cat*_ values of Mo- and V-Nase were varied to see at what point V-Nase growth rate exceeds that of Mo-Nase or Mo-Nase does the same compared to V-nase. At first, the *k*_*cat*_ of V-Nase was increased to a very high value, but the growth rate of V-Nase was still lower than the WT and Mo-Nase. It indicates that the sensitivity of V-Nase activity with respect to temperature is very low. This finding is consistent with previously published work on another gram negative bacteria, *Azotobacter chroococcum* (67). Later, the *k*_*cat*_ of Mo-Nase was decreased to a very low value, and at that low *k*_*cat*_, Mo-nase growth was actually lower than the V-Nase and higher than the Fe-Nase, which is similar to the finding from literature (45). Therefore, by tuning the *k*_*cat*_, ME-model was able to capture the experimentally observed temperature sensitivity of different nitrogenase enzyme. FIG 8b qualitatively summarized the effect of *k*_*cat*_ on growth of Mo-only and V-only strains of *R. palustris*.

### Conclusion

In this work, the first ever ME-model of *R. palustris* was developed. Growth rates predicted by the ME-model for different substrates closely matched with experimental growth rate data. The ME-model also predicted a diminishing carbon fixation at the theoretical maximum growth and subsequently malate dehydrogenase and glycerol-3 phosphate dehydrogenase as alternate electron sinks. Furthermore, the ME-model postulated electron transport through ferredoxin as a key regulatory feature to distribute reduced cofactor pools between carbon and nitrogen fixation pathways. Finally, ME-modeling framework successfully captured experimentally observed temperature regulation of different nitrogenase enzymes.

Going froward, this ME-model can be used as a powerful platform to further characterize different features of *R. palustris* metabolism. Specially characterizing a complete profile of environment specific isozyme expressions and optimal protein allocation. Furthermore, this ME-model can be used to design and fine-tune mutants of *R. palustris* for metabolic engineering purpose. One such application can be to produce PHB, a bioplastic precursor, which has potential to replace petroleum-based plastics. Under anaerobic-photoheterotrophic growth of *R. palustris*, PHB can work as an electron sink (7). Our previous effort (7) successfully established three design strategies to select the ideal lignin breakdown products (LBPs) for commercial PHB production from *R. palustris*. This ME-modeling framework can be further used to gain similar regulatory insights, as discussed in this paper, on how electrons are distributed in PHB producing pathways when different LBPs are used as substrates.

## MATERIALS AND METHODS

### ME-model of *R. palustris*

In addition to the metabolic reactions from the M-model, ME-model consists of translation and transcription reactions along with metabolic reactions (FIG 9). In order to model transcription and translation reaction, GPR association of each reaction is required. The initial GPR association was collected from literature (59). Later that GPR association was manually curated using the detail genome annotation from literature (2).

**FIG 9.**
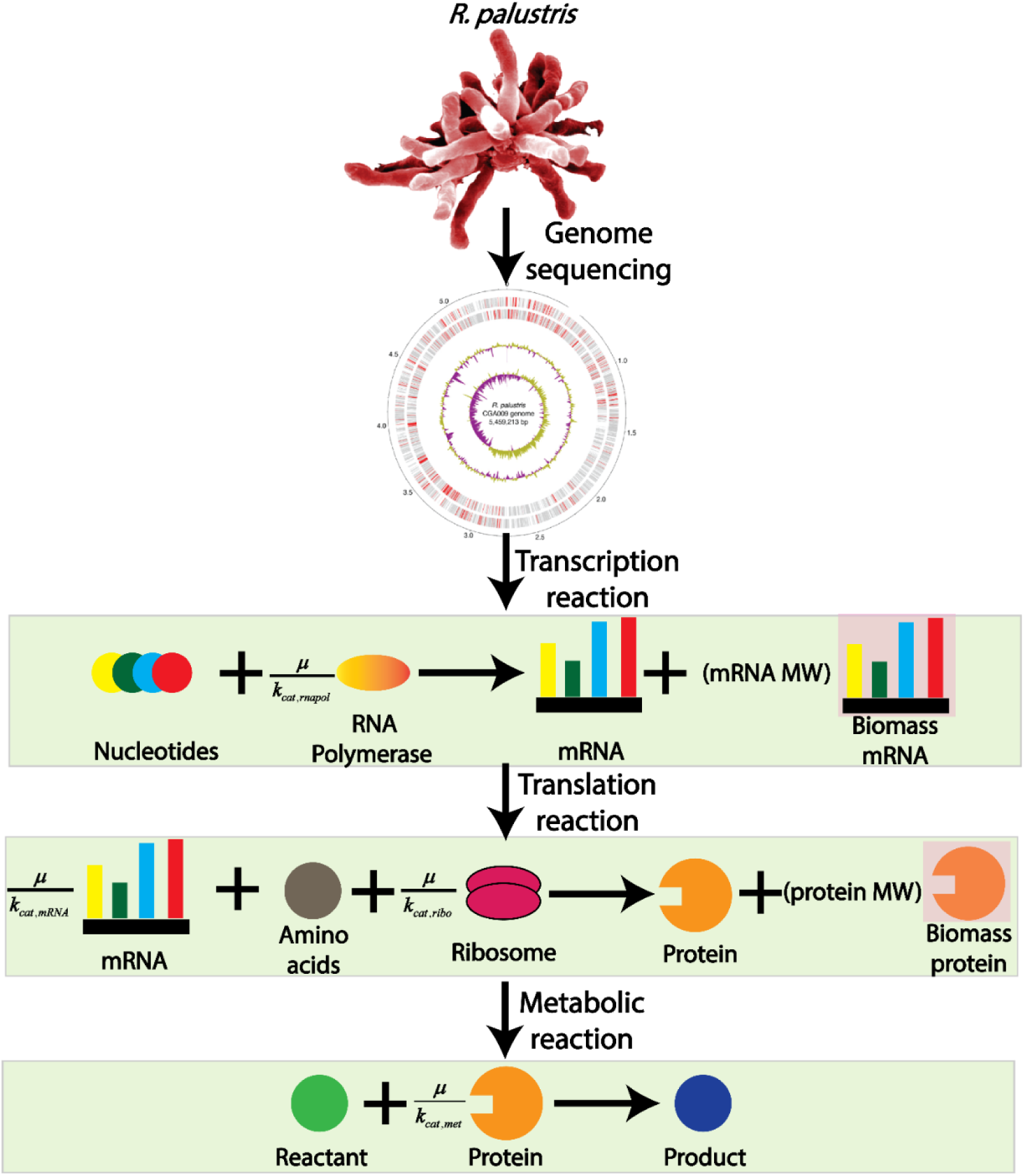
*R. palustris* ME-model reconstruction. In the M-model, only metabolic reactions are incorporated to perform genome-scale metabolic modeling. However, in the ME-modeling framework, transcription and translation process are also incorporated, adding two separate layer of regulation for metabolic reaction. Each layer of regulation are coupled with the biomass growth through catalytic turnover rate and the biomass growth. This process is known as coupling and the coupling parameter is in the form of 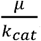, where *μ* indicates the growth rate and *k*_*cat*_ indicates the catalytic turnover rate for that.

This GPR association can be accessed in supplemental material Table 3. The overall ME-model reconstruction procedure was conducted in accordance to the COBRAme protocol (49) which is summarized in FIG 2. The ME-model is a multi-scale model; hence it requires the addition of coupling constraints to relate different cellular processes to each other. The coupling constraints are in the form of 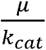. Here *μ* is the growth rate and *k*_*cat*_ approximates the effective turnover rate for the different macromolecules. Detailed mathematical description for 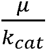 of different macromolecular process and values of different parameters can be found in the supplemental material Text S1 and in the original COBRAme protocol (49).

To calculate *k*_*cat*_ for different enzymes, a mean *k*_*cat*_ value of 65 *s*^−1^ was used, which was reported for the *E. coli* in another ME-modeling framework (31). This mean *k*_*cat*_ was modified for each enzyme based on the solvent accessible surface area (SASA), following the same ME-modeling framework (31). SASA can be defined as is the surface area of an enzyme that is accessible to a solvent. Also, a previous study (68) reported a correlation between SASA and molecular weight of the enzyme as following:

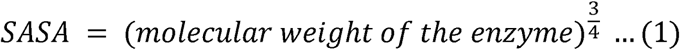

Overall, the following equation was used to calculate *k*_*cat,enzyme*_ for each enzyme, based on the mean turnover rate (*k*_*cat,mean*_), mean SASA (*sAsA*_*mean*_), and SASA for the specific enzyme (*sAsA*_*enzyme*_).

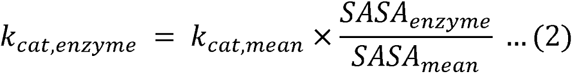

For transcription reactions, RNA polymerase is needed to produce the required mRNA for the protein production. RNA polymerase of *R. palustris* consists of five subunits: two alpha (*α*) subunits, a beta (*β*) subunit, a beta prime subunit (*α′*), and a small omega (*ω*) subunit (69). In the model each individual subunits were synthesized to form the RNA polymerase. Later, these RNA polymerase transform different nucleotide to mRNA.

For translation reactions, ribosomal RNA is required to transform amino acids into different proteins. *R. palustris* utilizes 70S ribosomes, each consisting of a small (30S) and a large (50S) subunit (70). The large subunit is composed of a 5S RNA subunit (120 nucleotides), a 23S RNA subunit (2900 nucleotides), and 31 proteins. The small subunit is composed of a 16S RNA subunit (1542 nucleotides) and 21 proteins (70). It was also assumed that tRNA charging of amino acid to the ribosome was not a rate limiting process in the translation reaction. Hence no macromolecular synthesis of tRNA was included in the model.

For each transcription or translation reaction in the ME-model, an amount of a biomass protein and biomass mRNA were produced with a stoichiometry equal to the molecular weight (in *kDA*) of the protein or mRNA being made. FIG 9 shows an example of this where the translation reaction produces both the catalytic protein as well as the biomass protein. Similarly, the transcription reaction produces mRNA required for the protein synthesis and also biomass mRNA requirements. The biomass protein and mRNA participate in the ME-model biomass dilution reaction, restricting the total biomass components production equal to the rate of biomass dilution.

Transcription and translation reactions were included for all reactions for which GPR are available. For remaining pathways, an enzyme was used with an average length of 283 amino acids and molecular weight of 31.09 *kDA*.

To capture the differential expression of the carbon fixing isozymes, a constraint was added to the ME-model to account for the co-expression of both rubisco form I and form II as follows:

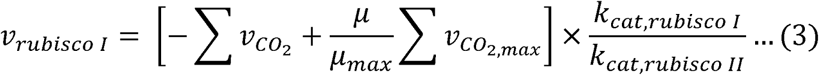

In equation (1), *v*_*rubisco I*_ represents the expression of rubisco form I, which is a function of carbon dioxide generation 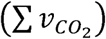, growth rate (*μ*), theoretical maximum growth rate (*μ*_*max*_), carbon dioxide generation at theoretical maximum growth rate 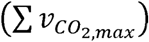, and effective catalytic rate of rubisco form I (*k*_*cat,rubisco I*_) and rubisco form II (*k*_*cat,rubisco II*_).

For each of the substrate, the total ATP production by the ME-model was capped according to the following equation proposed in the literature (7):

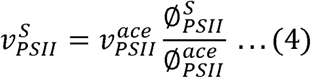

Here “S” and “ace” refer to different substrates and acetate, respectively. Also, 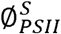 and 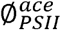 refer to the photosynthetic yield of different substrates and acetate respectively. Photosynthetic yields for different substrates are collected from literature (7).

### Accuracy calculation in the validation study

In the validation study, using the ME-model, aerobic growth of *R. palustris* was simulated with *p*-coumarate and succinate as sources of carbon and (*NH*_4_)_2_*so*_4_ as a sole source of nitrogen. From the ME-model, fluxes of transcriptomics and proteomics reactions were calculated for both carbon sources. Considering the transcriptomics and proteomics reaction fluxes for succinate uptake as the baseline condition, fold changes for all the gene expression and protein was calculated for *p*-coumarate uptake. If the fold change is greater than 1, it was noted as upregulated. If the fold change is less than 1, it was noted as downregulated. Once the upregulated/downregulated fold changes of transcription and translation reactions were calculated, that fold changes were compared with the literature (34). If both fold changes, from the ME-model and the experimental study, showed same direction (upregulated or downregulated) of fold change, then the prediction is correct. Otherwise the prediction is incorrect. Accuracy was then calculated as a percentage between correct prediction and total predictions.

### Simulation tools and software

The General Algebraic Modeling System (GAMS) version 24.7.4 with IBM CPLEX solver was used to run pFBA algorithm on the model. The algorithm was scripted in GAMS and then run on a Linux-based high-performance cluster computing system at the University of Nebraska-Lincoln.

## SUPPLEMENTARY MATERIALS

Supplemental material is available online only.

FIG S1-S4, PDF file, 0.01 MB.

TABLE S1, DOCX file, 1.7 MB

TABLE S2, DOCX file, 0.05 MB.

TABLE S3, DOCX file, 0.2 MB

TEXT S1, DOCX file, 0.03 MB

## DATA AVAILABILITY

All the codes used in this work can be found in the following GitHub directory: https://github.com/ssbio/palustris_ME_model

## ACKNOWLEDGEMENT

We gratefully acknowledge funding support from NSF Career Award grant number 1943310.

