## Supplementary material for "Characterizing the interplay of rubisco and nitrogenase enzymes in anaerobic-photoheterotrophically grown *Rhodopseudomonas palustris* CGA009 through a genome-scale metabolic and expression model": FIG S1-S4

Theoretical vs Experimental Growth

Theoretical growth rate (per day)

Experimetnal growth rate (per day)

| WIld type and other mutants | Theoretical maximum growth |
| --- | --- |
| WT | 1.378 |
| Mo-Nase | 1.378 |
| V-Nase | 1.221 |
| Fe-Nase | 0.912 |

|  | Experimental growth |
| --- | --- |
| WT | 0.7 |
| Mo-Nase | 0.65 |
| V-Nase | 0.57 |
| Fe-Nase | 0.45 |

**FIG S1** Comparison between theoretical maximum growth found from the ME-modeling framework and the experimental work from Luxem et al. 2020 when succinate was used as substrate.

Acetate WT

Growth, Nitrogenase flux

| Acetate uptake | Growth | Mo-nase |
| --- | --- | --- |
| 0 | 0 | 0 |
| 2 | 0.286 | 0.281 |
| 4 | 0.571 | 0.564 |
| 6 | 0.856 | 0.848 |
| 8 | 1.141 | 1.134 |
| 10.243 | 1.213 | 1.207 |

**FIG S2 (a)** For acetate uptake, growth rate and nitrogen fixation rate for WT *R. palustris.* For each of the case, growth rate and nitrogen fixation closely follow each other.

Acetate Mo-Nase

Growth, Nitrogenase flux

| Acetate uptake | Growth | Mo-nase |
| --- | --- | --- |
| 0 | 0 | 0 |
| 2 | 0.286 | 0.281 |
| 4 | 0.571 | 0.564 |
| 6 | 0.856 | 0.848 |
| 8 | 1.141 | 1.134 |
| 10.243 | 1.213 | 1.207 |

**FIG S2 (b)** For acetate uptake, growth rate and nitrogen fixation rate for Mo-Only mutant*.* For each of the case, growth rate and nitrogen fixation closely follow each other.

Acetate V-Nase

Growth, Nitrogenase flux

| Acetate uptake | Growth | Mo-nase |
| --- | --- | --- |
| 0 | 0 | 0 |
| 2 | 0.286 | 0.281 |
| 4 | 0.571 | 0.564 |
| 6 | 0.856 | 0.848 |
| 8 | 1.055 | 1.048 |
| 9.2 | 1.09 | 1.083 |

**FIG S2 (c)** For acetate uptake, growth rate and nitrogen fixation rate for V-Only mutant*.* For each of the case, growth rate and nitrogen fixation closely follow each other.

Acetate Fe-Nase

Growth, Nitrogenase flux

| Acetate uptake | Growth | Mo-nase |
| --- | --- | --- |
| 0 | 0 | 0 |
| 1 | 0.143 | 0.14 |
| 2 | 0.286 | 0.281 |
| 3 | 0.429 | 0.423 |
| 5 | 0.714 | 0.706 |
| 7.05 | 0.837 | 0.829 |

**FIG S2 (d)** For acetate uptake, growth rate and nitrogen fixation rate for Fe-Only mutant*.* For each of the case, growth rate and nitrogen fixation closely follow each other.

Theoretical vs Experimental Growth

Theoretical growth rate (per day)

Experimental growth rate (per day)

| Wild type and other mutants | Theoretical maximum growth |
| --- | --- |
| WT | 1.213 |
| Mo-Nase | 1.213 |
| V-Nase | 1.09 |
| Fe-Nase | 0.837 |

|  | Experimental growth |
| --- | --- |
| WT | 0.744167 |
| Mo-Nase | 0.59625 |
| V-Nase | 0.795 |
| Fe-Nase | 0.460556 |

(e)

**FIG S2 (e)** For acetate uptake, comparison between theoretical maximum growth found from the ME-modeling framework and the experimental work from Luxem et al. 2020 when acetate was used as substrate.

Butyrate WT

Growth, Nitrogenase flux

| Butyrate uptake | Growth | Mo-nase |
| --- | --- | --- |
| 0 | 0 | 0 |
| 1 | 0.286 | 0.281 |
| 2 | 0.571 | 0.564 |
| 3 | 0.856 | 0.848 |
| 4 | 1.141 | 1.134 |
| 5.497 | 1.29 | 1.285 |

**FIG S3 (a)** For butyrate uptake growth rate and nitrogen fixation rate for WT *R. palustris*. For each of the case, growth rate and nitrogen fixation closely follow each other.

Butyrate Mo-Nase

Growth, Nitrogenase flux

| Butyrate uptake | Growth | V-nase |
| --- | --- | --- |
| 0 | 0 | 0 |
| 1 | 0.286 | 0.281 |
| 2 | 0.571 | 0.564 |
| 3 | 0.856 | 0.848 |
| 4 | 1.141 | 1.134 |
| 5.497 | 1.29 | 1.285 |

**FIG S3 (b)** For butyrate uptake growth rate and nitrogen fixation rate for Mo-only mutant. . For each of the case, growth rate and nitrogen fixation closely follow each other.

Butyrate V-Nase

Growth, Nitrogenase flux

| Butyrate uptake | Growth | Fe-nase |
| --- | --- | --- |
| 0 | 0 | 0 |
| 1 | 0.286 | 0.281 |
| 2 | 0.571 | 0.564 |
| 3 | 0.856 | 0.848 |
| 4 | 1.094 | 1.087 |
| 4.9 | 1.152 | 1.145 |

**FIG S3 (c)** For butyrate uptake growth rate and nitrogen fixation rate for V-only mutant. . For each of the case, growth rate and nitrogen fixation closely follow each other.

Butyrate Fe-Nase

Growth, Nitrogenase flux

| Butyrate uptake | Growth | Mo-nase |
| --- | --- | --- |
| 0 | 0 | 0 |
| 1 | 0.286 | 0.281 |
| 2 | 0.571 | 0.564 |
| 2.5 | 0.714 | 0.706 |
| 3 | 0.835 | 0.827 |
| 3.69 | 0.873 | 0.865 |

**FIG S3 (d)** For butyrate uptake growth rate and nitrogen fixation rate for Fe-only mutant. For each of the case, growth rate and nitrogen fixation closely follow each other.

Theoretical vs Experimental Growth

Theoretical growth rate (per day)

Experimetnal growth rate (per day)

| WIld type and other mutants | Theoretical maximum growth |
| --- | --- |
| WT | 1.29 |
| Mo-Nase | 1.29 |
| V-Nase | 1.152 |
| Fe-Nase | 0.873 |

|  | Experimental growth |
| --- | --- |
| WT | 0.82 |
| Mo-Nase | 0.6 |
| V-Nase | 0.66 |
| Fe-Nase | 0.48 |

**FIG S3 (e)** For butyrate uptake, comparison between theoretical maximum growth found from the ME-modeling framework and the experimental work from Luxem et al. 2020 when acetate was used as substrate.

p-Coumarate WT

Growth, Nitrogenase flux

| p-Coumarate uptake | Growth | Mo-nase |
| --- | --- | --- |
| 0 | 0 | 0 |
| 0.5 | 0.321 | 0.316 |
| 1 | 0.643 | 0.636 |
| 1.5 | 0.963 | 0.955 |
| 2 | 1.143 | 1.136 |
| 2.562 | 1.213 | 1.207 |

**FIG S4 (a)** For *p*-Coumarate uptake growth rate and nitrogen fixation rate for (a) WT *R. palustris*. For each of the case, growth rate and nitrogen fixation closely follow each other

p-Coumarate Mo-Nase

Growth, Nitrogenase flux

| p-Coumarate uptake | Growth | Mo-nase |
| --- | --- | --- |
| 0 | 0 | 0 |
| 0.5 | 0.321 | 0.316 |
| 1 | 0.643 | 0.636 |
| 1.5 | 0.963 | 0.955 |
| 2 | 1.143 | 1.136 |
| 2.562 | 1.213 | 1.207 |

**FIG S4 (b)** For *p*-Coumarate uptake growth rate and nitrogen fixation rate for Mo-only mutant. For each of the case, growth rate and nitrogen fixation closely follow each other.

p-Coumarate V-Nase

Growth, Nitrogenase flux

| p-Coumarate uptake | Growth | Fe-nase |
| --- | --- | --- |
| 0 | 0 | 0 |
| 0.5 | 0.321 | 0.316 |
| 1 | 0.643 | 0.636 |
| 1.5 | 0.963 | 0.955 |
| 2 | 1.055 | 1.048 |
| 2.3 | 1.09 | 1.078 |

**FIG S4 (c)** For *p*-Coumarate uptake growth rate and nitrogen fixation rate for V-only mutant. For each of the case, growth rate and nitrogen fixation closely follow each other

p-Coumarate Fe-Nase

Growth, Nitrogenase flux

| p-Coumarate uptake | Growth | Fe-nase |
| --- | --- | --- |
| 0 | 0 | 0 |
| 0.25 | 0.161 | 0.158 |
| 0.5 | 0.321 | 0.316 |
| 1 | 0.643 | 0.636 |
| 1.5 | 0.812 | 0.804 |
| 1.7 | 0.837 | 0.829 |

**FIG S4 (d)** For *p*-Coumarate uptake growth rate and nitrogen fixation rate for Fe-only mutant. For each of the case, growth rate and nitrogen fixation closely follow each other.

Theoretical vs Experimental Growth

Theoretical growth rate (per day)

| Wild type and other mutants | Theoretical maximum growth |
| --- | --- |
| WT | 1.213 |
| Mo-Nase | 1.213 |
| V-Nase | 1.09 |
| Fe-Nase | 0.837 |

**FIG S4 (e)** For *p*-Coumarate uptake, theoretical maximum growth found from the ME-modeling framework for WT *R. palustris* and other -only mutants.
