## Supplementary material for "Characterizing the interplay of rubisco and nitrogenase enzymes in anaerobic-photoheterotrophically grown *Rhodopseudomonas palustris* CGA009 through a genome-scale metabolic and expression model": Text S1

Transcription Reaction:

Coefficient of ribosome consumption = $\frac{l_{TU,i}}{3c_{ribo}K_{T}}\left( \mu+r_{0}K_{T} \right)$

Here,

$l_{TU,i} =$Number of nucleotides in each mRNA.

$c_{ribo} = \frac{m_{rr}}{m_{aa}f_{rRNA}}$, where $m_{rr}$ is mass of rRNA per ribosome, $m_{aa}$ is molecular weight of average amino acid, and $f_{rRNA}$is the fraction of RNA that is rRNA. The value of these parameters were chosen as following from the literature (1):

$m_{rr} = 1700 KDa$, $m_{aa} = 109 Da$, and $f_{rRNA} = 0.86$

$\mu=$ Growth rate.

$r_{0} =$The value of this dimensionless constant was set to 4.5 (1).

$K_{T} =$This is defined as the effective catalytic constant which connects the RNA-to-Protein ratio with the growth rate. The value of this parameter was set to $4.5 h^{-1}$ (1).

Translation Reaction:

Coefficient of mRNA consumption = $\frac{\mu+r_{0}K_{T}}{{3c}_{mRNA}K_{T}}$

Here,

$c_{mRNA} =$ $\frac{m_{nt}}{m_{aa}f_{mRNA}}$, where $m_{nt}$ is the average molecular mass of RNA nucleotide, and $f_{mRNA}$ is the fraction of RNA that is mRNA. The value of these parameters were chosen as following from the literature (1):

$m_{nt} =324 Da$, and $f_{rRNA} = 0.02$

Coefficient of amino acid consumption = $\frac{l_{p,i}}{c_{ribo}K_{T}}\left( \mu+r_{0}K_{T} \right)$

Here,

$l_{p,i} =$ Number of amino acids in each proteins.

Metabolic Reaction:

Coefficient of mRNA consumption = $\frac{\mu}{k_{cat}}$

Here,

$k_{cat} =$ Effective turnover rate. The mean effective turnover rate used in this study is $65 s^{-1}$, which was reported for the *E. coli* in another ME-modeling framework (1). For different enzymes, this mean effective turnover rate was modified based on the molecular weight of the enzyme.

Metabolite pool size calculation:

Let’s assume there are thee reactions as following”

Reaction 1: $C\to A$

Reaction 2: $D\to3A$

Reaction 3: $A\to4E$

Here, in reaction 1 and 2, A is being produced and in reaction 3, A is being consumed. If the flux of reaction 1 is $v_{1}=1 \frac{mmol}{gDW.day}$ and the flux of reaction 2 is $v_{2}=3 \frac{mmol}{gDW.day}$ then applying the pseudo steady state principles, the flux of reaction 3 will be 10 $\frac{mmol}{gDW.day}$. Then, the overall pool size of A will be $v_{3}=10 \frac{mmol}{gDW.day}$.

The idea is demonstrated in the following figure.

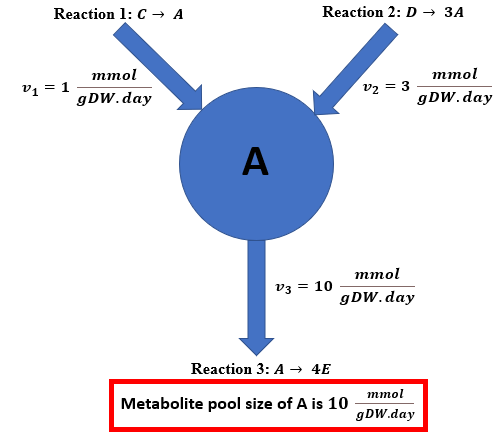

**FIG S5** Demonstrating the calculation of metabolite pool size. Here A is a fictitious metabolite that is produced from two reactions, 1 and 2. Reaction 1 has a flux of 1 $\frac{mmol}{gDW.day}$ and reaction 2 has a flux of 3 $\frac{mmol}{gDW.day}$. A is only consumed in reaction 3. Applying pseudo steady mass balance assumption, flux of reaction 3 will be 10 $\frac{mmol}{gDW.day}$. Hence, the metabolite pool size of A is 10 $\frac{mmol}{gDW.day}$.
